# Stimulus contrast information modulates sensorimotor decision making in goldfish

**DOI:** 10.1101/849521

**Authors:** Santiago Otero Coronel, Nicolás Martorell, Martín Berón de Astrada, Violeta Medan

## Abstract

Animal survival relays on environmental information gathered by their sensory systems. We found that contrast information of a looming stimulus biases the type of defensive behavior goldfish (*Carassius auratus*) perform. Low contrast looms only evoke subtle alarm reactions whose probability is independent of contrast. As looming contrast increases, the probability of eliciting a fast escape maneuver, the C-start response, increases dramatically. Contrast information also modulates the decision of when to escape. Although looming retinal size is determinant of response latency, we found that contrast acts as an additional parameter influencing this decision. When presented progressively higher contrast stimuli, animals need shorter periods of stimulus integration to initiate the response. Our results comply the notion that the decision to escape is a flexible process initiated with stimulus detection and followed by assessment of the perceived risk posed by the stimulus. Highly disruptive behaviors as the C-start are only observed when a multifactorial threshold that includes stimulus contrast is surpassed.

**Summary statement:** This study highlights that in fish, the decision of what to do after threat detection is a multifactorial non-binary process that includes assessing the relative contrast of the potential threat. Increasingly higher contrast stimuli produce a progressive increase in C-start escape probability and a decrease in response latency. More subtle alarm reactions are, on the contrary, mostly insensitive to changes in contrast. This might reflect that while subtle reactions have lower thresholds to be executed, disruptive behaviors as the C-start must surpass higher saliency thresholds that integrate multiple aspects including contrast.

## INTRODUCTION

Evasive behaviors are essential to avoid harm from predators or other threats in the environment. Although critical for animal survival, escaping comes at the cost interrupting other behaviors such as foraging or mating, and thus it is not performed unless the perceived threat surpasses a decision threshold. To match behavior to perceived risk animals first detect and then evaluate threat levels to decide to perform (or not) an escape. Perceived threat levels depend not only on the characteristics of the stimulus but also on the internal state of the animals and previous experience (Evans et al., 2019).

One of the best studied escape behaviors is the C-start of fish (Batty, 1989; Dill, 1974; Eaton et al., 1991; Faber et al., 1989; Kohashi and Oda, 2008; Neumeister et al., 2010; Preuss and Faber, 2003; Whitaker et al., 2011). The C-start is a high threshold escape behavior consisting on a first stage where fast and massive unilateral contraction of trunk muscles results in the fish adopting a C-shape followed by a return stroke in the opposite direction (‘return flip’) where the tail straightens propelling the animal away from the potential danger (Domenici and Blake, 1997; Eaton et al., 1977; Zottoli, 1977). Although the initial stage is highly stereotyped and its directionality mostly imposed by the direction of the threat, it can be modulated by the presence of obstacles or other fish (Domenici, 2010; Eaton and Emberley, 1991).

In laboratory conditions, robust C-start behavior can be elicited by visual looming threats (Dunn et al., 2016; Preuss et al., 2006; Temizer et al., 2015). Looming stimuli usually consist on a computer-generated black disks rapidly expanding over a white background. These types of stimuli have been shown to induce escape behaviors from invertebrates to humans (Laurent and Gabbiani, 1998) suggesting that the neural circuits involved in avoiding an approaching predator or a collision have evolved early during evolution (Evans et al., 2019). Fish can compute looming velocity and retinal angular size to decide when to initiate a C-start (Dunn et al., 2016; Heap et al., 2018; Preuss et al., 2006; Temizer et al., 2015) and to adjust the kinematics of the escape swim (Bhattacharyya et al. 2017). However, before deciding to execute a C-start, fish have to evaluate threat levels, a necessary previous step which has not been studied in detail.

The level of perceived risk posed by a stimulus will depend on its specific characteristics as well as internal state and prior experience of the animal. Escape thresholds can rise when animals are feeding or when previous encounters with the stimulus had no harmful consequences (Lima and Dill, 1990; Lloyd and Dayan, 2018; Roberts et al., 2019, 2016). Vigilance levels can also affect threat detection (De Franceschi et al., 2016). For example, in fish that are actively exploring the environment, threat detection can produce an interruption of ongoing locomotion to stabilize the visual panorama to facilitate tracking the stimulus.

Here we varied the contrast of a looming stimuli to manipulate its salience and investigated the effect on the behavioral choices fish performed. In addition, we tested the hypothesis that stimulus contrast is incorporated in the computing mechanism that decides response latency.

## METHODS

### Animals

Adult goldfish (*Carassius auratus*) of both sexes, 7–10 cm of standard body length, were purchased from FunFish (Córdoba, Argentina). Fish were allowed to acclimate for at least a week after transport and were kept in rectangular glass holding tanks (30×60×30 cm; 95 l) in groups of 10 animals. Tanks were supplied with filtered and dechlorinated water and maintained at 18 °C. Ambient light was set to a 12 hr light/dark photoperiod. Animals were fed floating pellets (Sera, Germany) five times a week.

All animal procedures were performed in accordance with the guidelines and regulations of the Institutional Animal Care and Use Committee of Facultad de Ciencias Exactas y Naturales, Universidad de Buenos Aires (protocol #70).

### Experimental set-up and behavioral protocol

Goldfish were tested in a rectangular experimental tank (48 cm length, 36 cm width and 27 cm height) with its external walls covered with white opaque cardboard to avoid external visual stimulation. In addition, opaque panels covered all sides and top of the experimental setup, preventing external light to reach the tank. Experiments were made in a silent room with ceiling lights off. The experimental tank was filled with filtered dechlorinated water up to a height of 20 cm. A liquid crystal display (LCD) screen used for visual stimulation was secured 6 cm above the water surface (Fig. 1A). The long axis of the screen was placed parallel to the long axis of the tank. Illumination was homogeneous in all the tank and no shelters were provided. The tank was situated on a transparent acrylic sheet allowing video recording of the fish’ behavior and stimulus presentation from beneath at 240 or 480 fps (Casio EX ZR100).

**Figure 1.**
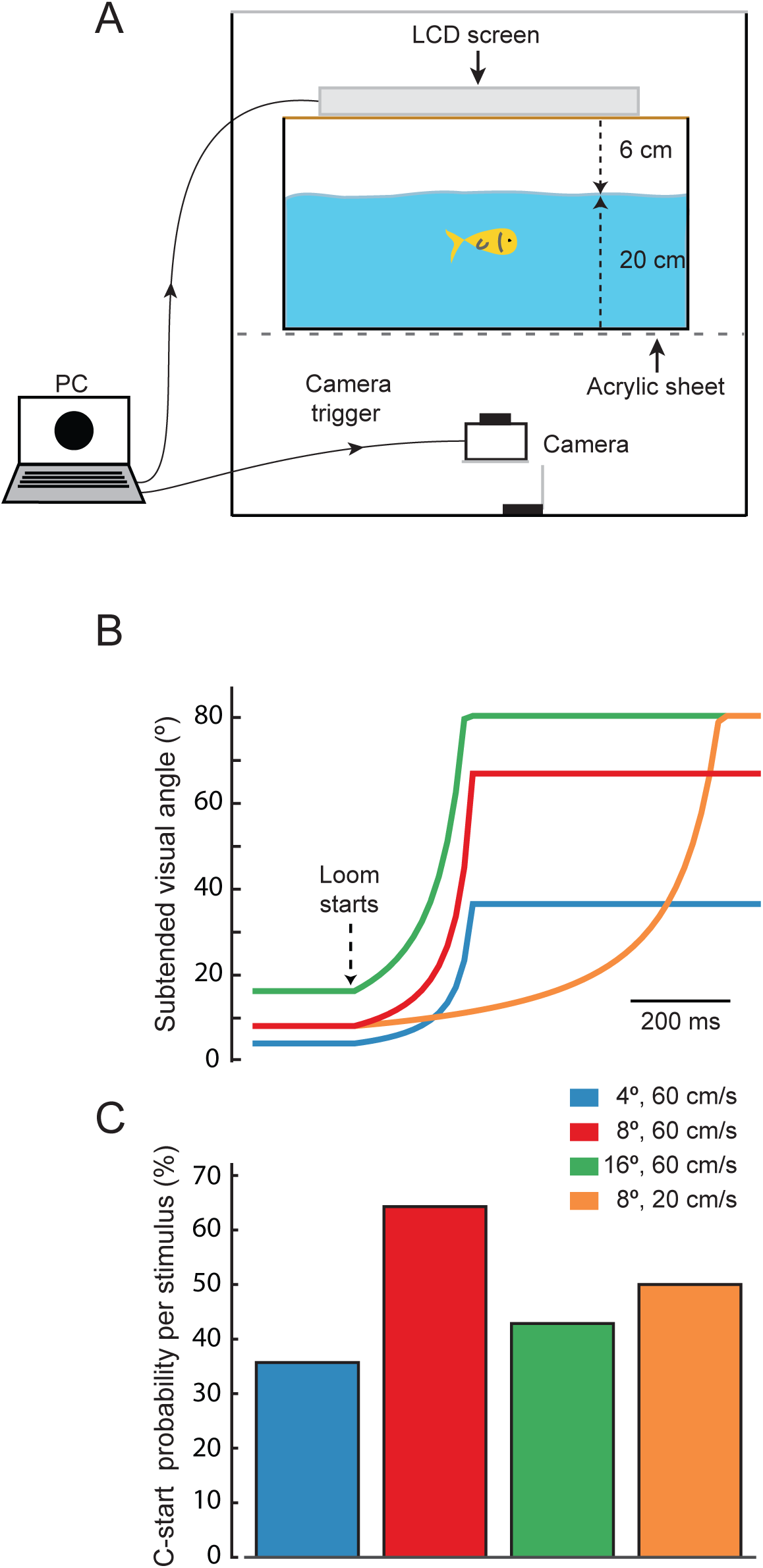
Looming evoked escapes in goldfish. A. Behavioral setup. Computer-generated visual looms were delivered through an LCD screen located on top of the experimental aquarium which stands on a transparent acrylic platform. Fish behavior and looming expansion was filmed from below with a camera acquiring at 240 or 480 fps. The same computer triggered acquisition of the camera. B. Time course of the stimuli used to characterize response probability to high contrast visual looms. The four stimuli differed in their initial subtended angle (4°, 8° or 16°) or expansion velocity (20 or 60 cm/s). C. C-start probability for each of the looms used. After this initial characterization the stimulus that evoked maximal C-start probability, loom onset at 8° and expanding at 60 cm/s (red), was chosen for the rest of the experiments.

Computer controlled presentation of visual stimuli on the LCD screen and triggering of the camera acquisition occurred 1.3 s before the stimulus appeared and stopped at 9.7 s after the end of visual stimulation. In addition, a small web camera recorded fish activity (60 fps) from below and allowed to monitor animal activity during the experiment.

Individual fish were placed in the experimental tank and allowed to acclimate for 30 minutes. Unless otherwise stated, the animal was then stimulated three times with the same looming stimulus with a 5-minute interval between presentations. After the experiment, the animal was returned to its holding tank.

### Visual stimuli

Computer generated black disks that expand over a white background (loom) efficiently elicit C-start escapes (Dunn et al., 2016; Medan and Preuss, 2014; Preuss et al., 2006; Temizer et al., 2015). In our study, looms were presented using a 380×305 mm LCD (1280×1024 pixels, refresh rate 75Hz, HP L1940T, Hewlett-Packard, USA). We changed the RGB value of the disk stimulus while keeping the background white to obtain various intensity contrast stimuli (IC).

To express intensity contrast we adopted the Michelson index (%), where contrast is defined as (I_loom_ − I_bkgn_) × 100/ (I_loom_ + I_bkgn_) and I_loom_ and I_bkgn_ refer to the irradiance of the expanding disk and the background respectively. The background had the maximum intensity the LCD can deliver in grayscale (i.e. the same three values in the RGB code; value 255; I_bkgn_ = 221.6 mW/m^2^) and the disk was set to different shades of grey (grayscale value 1-254) or black (grayscale value 0) according to the experimental group. The contrast of the different intensity contrast looms (IC) will be subsequently denoted by their subindex (e.g. IC_1.7_ represents an IC with a Michelson index of 1.7%). All light irradiance measurements were done with the irradiance sensor (J1812) of a Tektronix J17 photometer (Wilsonville, Oregon, USA) positioned in the center of the tank facing the LCD.

Escape responsiveness in fish depends on the dynamics of the visual loom stimulus (Bhattacharyya et al., 2017; Dunn et al., 2016; Preuss et al., 2006; Temizer et al., 2015), thus, an ineffective loom dynamic could represent a confounding factor when trying to detect contrast sensitivity with a looming stimulus. To ensure that our stimulus dynamic was efficiently triggering escape responses, we tested ICS in which black disks expanded over a white background with four different dynamics (Michelson contrast 89%, Fig. 1B). To calculate the angular retinal size of the disk, we assumed the fish to be in the center of the tank at the midpoint of the water column (Fig. 1A). Three of the four stimuli simulated the approximation of a sphere at 60 cm/s that subtended an angle of 4°, 8° or 16° at its stationary initial position and expanded up to 37°, 67° or 80° respectively in 221 ms (Fig. 1B). The fourth stimulus simulated a sphere moving at 20 cm/s that initially subtended an angle of 8° and reached 80° in 731 ms. Each of the four loom dynamics was tested twice in random order on each animal with an intertrial interval of 5 min (Fig. 1C). We found that although all stimuli were effective, the stimulus that subtended an initial angle of 8° and a velocity of 60 cm/s provoked the C-start response with highest probability (64%, N=7, n=14, Fig. 1C, red bar). We therefore used this dynamic (initial angle of 8° and a velocity of 60 cm/s) for the rest of the experiments.

### Data analysis

#### Behavioral responses

##### C-start escape responses

Videos were analyzed off line using ImageJ (NIH, Bethesda). Visual inspection of the videos allowed us to confirm the manual scoring of occurrence of C-start escape responses observed during the experiment and to measure its latency with ±2 ms error for videos recorded at 480 fps and ±4 ms error for videos recorded at 240 fps. The first frame at which the expanding loom attained its maximum size was considered as 0 ms. Therefore, C-start responses occurring before the end of the expansion have a negative latency while those occurring after the end of the expansion rendered a positive latency.

##### Alarm responses

Videos were also inspected and scored by three independent experimenters to analyze the occurrence of behaviors other than C-start responses. These responses included behaviors suggesting increased arousal and alarm. Alarm responses consist on a variety of subtle but robust motor reactions including accelerating or decelerating swimming, darting (a single fast acceleration in one direction with the use of the caudal fin); erratic movements/zig-zagging (representing fast acceleration bouts in rapid succession); and rapid abduction of fins with no body displacement (Kalueff et al., 2013; Laming and Savage, 1980; Savage, 1971). An alarm response was computed when all three observers agreed on the occurrence and description of the behavior.

### Statistical analysis

R Studio (version 3.5.0) was used for statistical analysis. A significance level of α= 0.05 was used throughout the study. The effect of looming contrast on the probability of executing alarm responses or an escape behavior was assessed with a binomial generalized linear model (GLM) considering contrast levels as a fixed factor. The effect of varying contrast on latency was analyzed with a linear model. Normality of the latency data was assessed using Shapiro’s test. Sample size is denoted by *N* when it refers to the animals used or *n* when it refers to the number of trials performed with each contrast level.

## RESULTS

As behavioral decisions are influenced by the immediate behavioral past, we first analyzed what the animals were doing 2 seconds before the expansion of loomings. As we were interested in behavioral decisions taken after detecting the stimulus, this analysis only includes IC stimuli that evoked behavioral responses in at least 30% of the animals, which was observed for ICs ranging from 89 to 2.8% (n=197, N=68). We classified the prior motor state of fish in three categories: 1) still, referring to animals that were only moving the pectoral fins with no net displacement of the body, 2) freezing, when we could not detect any movement other than occasional breathing movements and 3) swimming, when fish were actively moving the caudal fin and producing a net propulsion of their body. We next analyzed the transitions from those three pre-stimulation states to the different behavioral outcomes of the looming stimulation. The behaviors observed after looming presentation included those mentioned before and, in addition, incorporate C-start and alarm responses. Alarm responses were defined as sudden changes in swimming velocity and fin waving (see Methods section). Although freezing might be considered an alarm response (Kalueff et al. 2013) we treated it as a different category since, in contrast with the rest of alarm responses, it implied complete cessation of body movement. The Sankey diagram of Figure 2 depicts the proportion of previous motor state on the left and the behavioral responses observed after stimulation on the right. The diagram shows that 45% of all stimulations evoked a C-start. Two thirds of those responses (64%) were produced by animals that were either still (48%) or freezing (16%) before the expansion and only 36% of the responses correspond to animals that were swimming prior to the expansion. This suggests that being still might aid stabilizing visual panorama and therefore improve threat detection. The results also show that freezing does not preclude by itself the execution of an explosive response as the C-start as 56% of animals that were freezing responded with a C-start while the remaining 44% continued freezing. However, we never observed a transition from freezing to an alarm response.

**Figure 2.**
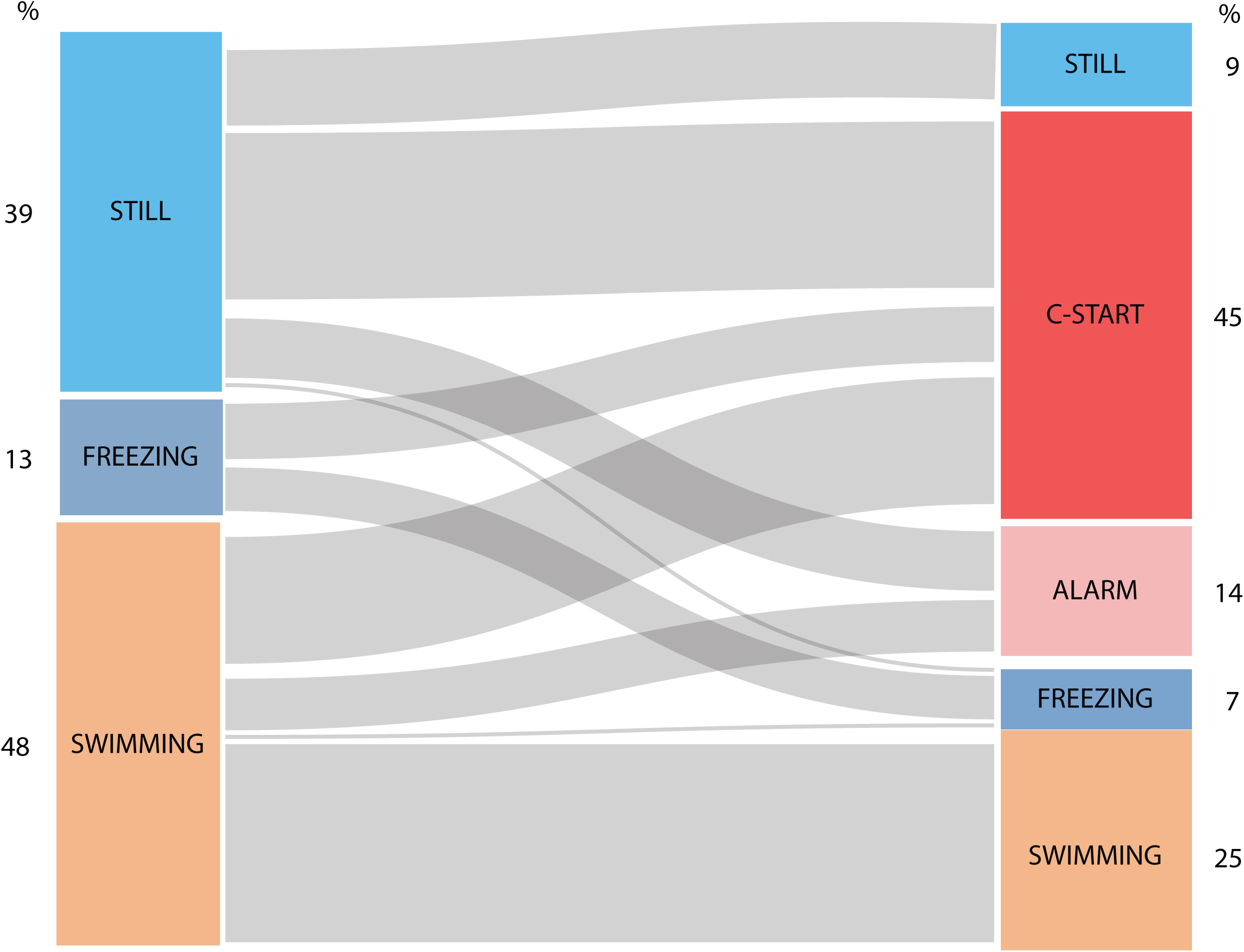
Sankey diagram of motor behavior before and after looming stimulation. The boxes on the left indicate the motor state before stimulation and the boxes on the right indicate the motor state after looming stimulation. The width of the grey lines are proportional to the numbers of animals transitioning from one type of behavior to other. Numbers on the outer edge of the boxes indicate the proportion of animals in each category.

In each of the three pre-stimulation behavioral categories defined, a proportion of animals did not modify their previous behavior. This could be either the result of animals detecting the stimulus and deciding not to alter their behavior or simply failure to detect the stimulus. In particular, animals that were freezing could have detected the stimulus and judged it threatening but deciding freezing to be the best response. Actively swimming animals, which, in the majority of cases (52%) did not change their behavior could have similarly failed to detect the stimulus. Alternatively, they may have judged interruption inadequate as they were already engaged in another activity (e.g. exploring the arena for a shelter or an exit). Our video analysis is insufficient to distinguish between these options.

After an evaluation period and depending on the risk perceived, fish will ignore the stimulus if no risk is detected, perform an alarm reaction if the perceived danger is intermediate or opt for a last-resource evasive behavior when danger is extreme. To specifically test if increasing the saliency of the stimulus modulates this decision making, we stimulated fish with 7 different looms of identical expansion dynamics but with contrasts that ranged from 0.7% to 89% Michelson Contrast (Figure 3). We found that although all looming stimuli had identical duration and identical subtended angle, increasing contrast produced a gradual switch from no evident motor reaction for looms of IC_0.7_ (83% did not alter their behavior, n=36) to an almost exclusive election of C-start escape for IC_89_ (90% performed C-starts, n=49; binomial GLM on effect of contrast on C-start probability, p<0.001). Curiously, alarm responses show no modulation by contrast as they were equally probable (between 11-20%) for all ICS with the exception of IC_89_ where only 4% of fish produced an alarm response (binomial GLM on the effect of contrast on the proportion of alarm responses, p=0.03).

**Figure 3.**
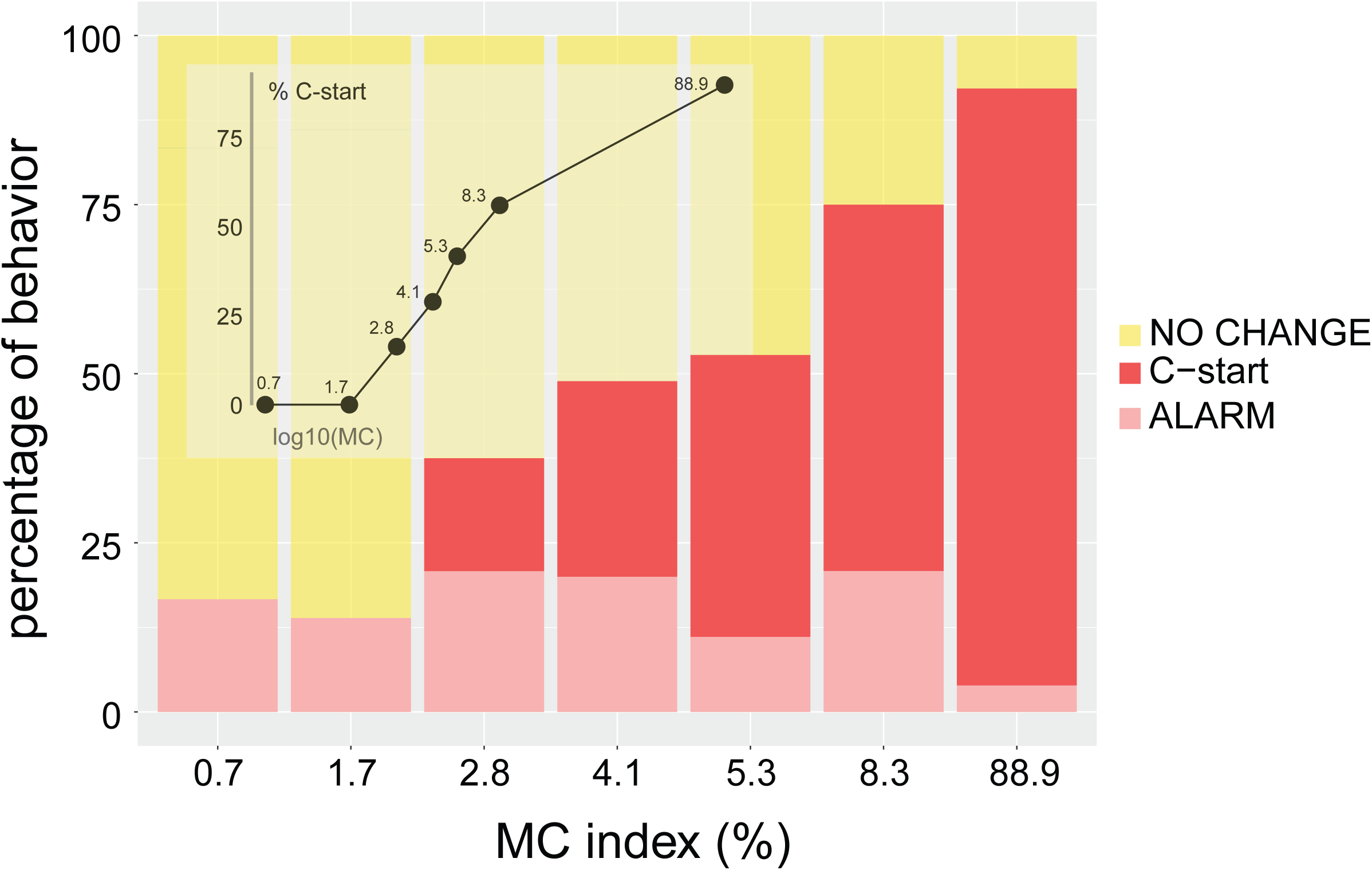
Increasing contrast shifts fish response to C-start behavior. The graph shows stacked bars indicating the relative proportion of C-starts (red), alarm responses (pink) or no change (yellow) in motor behavior. Inset shows the % C-start probability vs. MC represented in a logarithmic scale.

It has been repeatedly reported that black over white looms are more effective than white over black looms (Dunn et al., 2016; Preuss, 2006; Temizer et al., 2015). However, this is, to our best knowledge, the first evidence that goldfish can do more than to assign a binary (positive or negative) value to an expanding stimulus. Animals seem to attribute graded value to loom contrast when computing the threatening level of the loom. Inset of figure 3 shows the dependence between C-start probability and loom contrast. Threshold contrast for C-starts lies between IC_1.7_ and IC_2.8_ and then follows the Weber-Fechner law, as evidenced by the linear increase when the Michelson Contrast is represented on a logarithmic scale.

Since all the stimuli used in our experiments had the same expansion dynamics, we expected a fixed C-start latency (Bhattacharyya et al., 2017). Surprisingly, we found that goldfish progressively delay their escapes when saliency is lowered, i.e. they take progressively more time to initiate the C-start (Figure 4, linear model on the effect of contrast on latency, p<0.001, Shapiro test NS). While the highest contrast loom (IC_89_) produced mean (±SD) latencies of 69 ± 38 ms before end of the loom expansion, lower contrast looms delayed response more than 50 ms (IC_2.8_: 7±45 ms; IC_4.1_: 25±46 ms before end of the expansion). In fact, lower saliency looms evoked a significant proportion of C-starts that were so delayed that occurred after the end of the expansion (43% for IC_2.8_).

**Figure 4.**
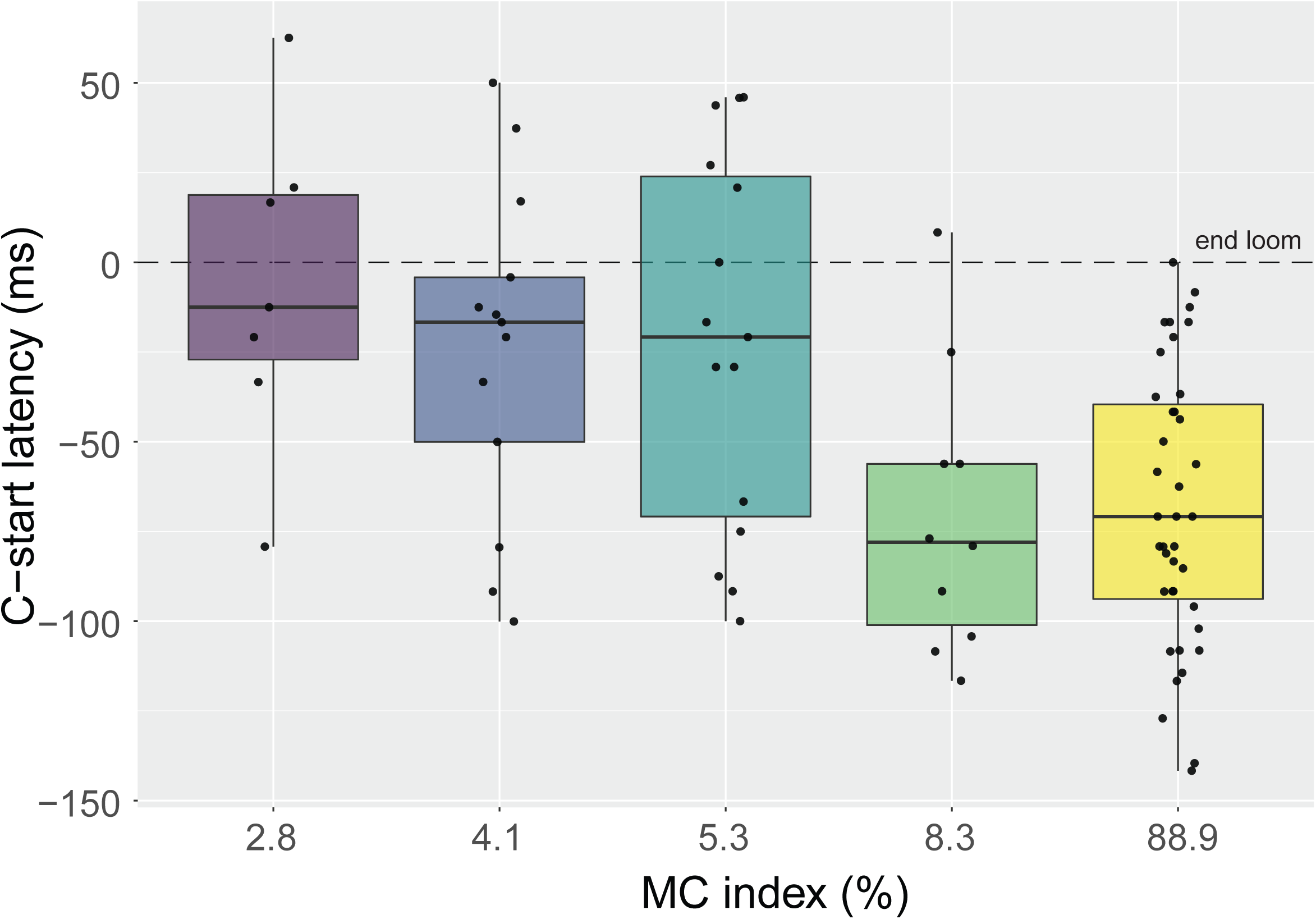
Stimulus saliency is correlated with C-start response latency. Looming stimuli of increasing Michelson index were associated with shorter response latencies. Boxplot represent median and 25th and 75 quartiles, and minimum and maximum values for each MC. Superimposed dots represent individual responses.

Since reducing the contrast of the stimuli increased the C-start response latency, we analyzed if other parameters of the response were also affected. We thus compared the kinematics of C-starts evoked by high and low IC (IC_89,_ IC_2.8_ and IC_4.1_) corresponding to 56 trails performed in 30 animals. If low and high contrast looms were recruiting the same sensorimotor networks we expected the main characteristics of stage 1 of the C-start to be similar. Indeed, animals startling to low or high contrast looms performed C-starts of comparable duration (median, IC_89_: 25ms; IC_4.1_: 21ms, IC_2.8_: 25ms, Fig. 5, upper panel) and angular bend (median, IC_89_: 71°; IC_4.1_: 70°, IC_2.8_: 70°, Fig. 5, lower panel) for both stimuli. Although these results do not exclude the possibility that other reticulospinal neurons are implicated, these results suggest that C-starts evoked by high or low contrast looms are conveyed through the Mauthner cell pathway.

**Figure 5.**
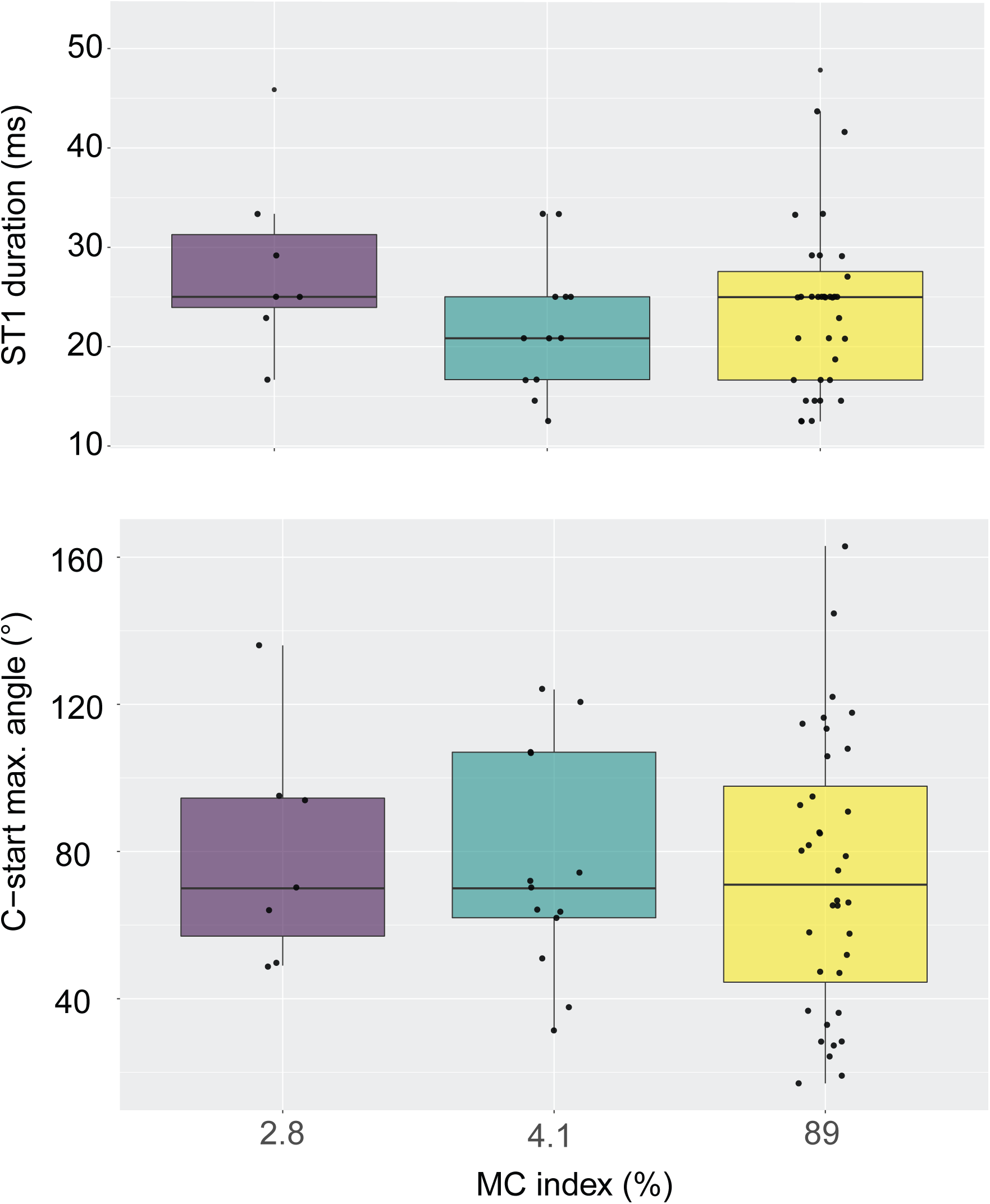
Stimulus saliency does not change C-start kinematics. Stage 1 duration (upper panel) or maximum stage 1 angle (lower panel) are similar for IC2.8, IC4.1 and IC89. Boxplot represent median and 25th and 75 quartiles, and minimum and maximum values for each MC. Superimposed dots represent individual responses.

## DISCUSSION

Although fast defensive behavior such as startle or escape responses were initially analyzed like reflex-like behaviors organized around command neurons (Eaton et al., 2001; Edwards et al., 1999) further investigation made it increasingly clear that defensive behaviors have a deeper complexity and flexibility that previously acknowledged (Card, 2012; Domenici, 2010; Evans et al., 2019; Simmons et al., 2010).

Studies on defensive responses in fish have mainly centered on the C-start escape response and the parameters of the auditory or visual stimuli that triggers this response. Our experiments revealed a wider behavioral flexibility of the defense response at two levels. First, we found that fish that detected the stimulus can display a range of behaviors that include but are not restricted to C-starts. Second, we found that if a C-start is performed, the time the animal takes to initiate it is modulated by the contrast of the stimulus.

When we varied the contrast of an otherwise invariant looming stimulus we observed that C-start probability decreased until disappearing for contrasts lower than IC2.8. Simultaneously, we observed a rather constant rate of alarm behaviors that only decreased for the highest contrast. This suggests that the initial process of stimulus detection seems to be, above a minimum sensitivity threshold, contrast independent. Although we surveyed a wide range of contrasts, we also found that the proportion of alarm responses were similar for all conditions excluding the highest contrast. However, we cannot rule out that contrast affects alarm behaviors. These are more subtle motor behaviors and thus we might have underestimated their frequency since we only included those events where scores by three independent observers matched. It could be possible that a reduction on alarm responses when contrast is reduced might be only detectable if more sensitive techniques are implemented. Heart rate frequency or skin conductance response levels are traditionally used to measure “alarm” (Burnovicz et al., 2009; Kreibig, 2010; Yoshida et al., 2009) although measuring analogous signals in freely swimming fish could be challenging.

In addition to modulating C-start probability, looming contrast also modulates *when* animals initiate the C-start. C-start response probability has been shown to be modulated by external factors such as spatial and social context (Eaton and Emberley, 1991; Fischer et al., 2015) and internal state variables such as hunger level or reproductive or social status (Filosa et al., 2016; Neumeister et al., 2010; Park et al., 2018). Furthermore, intrinsic characteristics of the stimulus such as its modality, temporal dynamics, or directionality have also been shown to shape the characteristics of the C-start. Specifically for visual loomings, the relationship between C-start latency and the loom velocity and subtended angle at the retina has been extensively studied (Bhattacharyya et al., 2017; Dunn et al., 2016; Preuss et al., 2006; Temizer et al., 2015). On the other hand, the dependency of the C-start with the contrast of the stimulus has only been previously investigated by presenting black over white or white over black combinations (Dunn et al., 2016; Medan and Preuss, 2014; Randlett et al., 2019; Temizer et al., 2015). In nature, however, there is a continuous scale of contrast between objects and background that animals could detect and extract information from. Here we found that, indeed, animals take into account the contrast to assign salience to a looming stimulus. Changing contrast of an otherwise identical loom stimuli is enough to modulate C-start probability and latency. If contrast is interpreted as a source of information, then high contrast looms provide enough information to reach decision threshold in a shorter time. As looming contrast diminishes, animals may need to integrate visual information for longer periods of time before reaching a C-start decision threshold, producing the increase in latency we observe.

Heap and colleagues (Heap et al., 2018) have recently proposed that visual information is conveyed through the retina not only to the optic tectum but also to the thalamus. They observed that thalamo-tectal projection neurons modulate the responses of looming sensitive tectal neurons. Luminance information carried by thalamic projection neurons increased C-start response rate and was found necessary to evoke directional escapes. The authors were capable of splitting luminance and expansion dynamics information of a looming stimulus. When they “reconstructed” the stimulus by simultaneously presenting the two cues to each eye independently they found a response enhancement, suggesting that contrast itself is capable of modulate response rate.

While that study did not vary luminance contrast, it would be reasonable to expect that the higher the luminance contrast the stronger the thalamic input to the tectum will be. This in turn would possibly lead to more intense input to downstream reticulospinal networks resulting in escapes with shorter latency. If true, this rationale would provide a mechanistic basis to the correlation we obtained between contrast and response latency.

The behavioral flexibility observed for less salient stimuli has been attributed to activity in different sets of tectal neurons which in turn innervate different populations of spinal projecting nuclei (Bhattacharyya et al., 2017). This is paralleled by the flexibility described for C-start escapes which depend on the activation of different reticulospinal neurons that in conjunction with the Mauthner cell are collectively known as the brainstem escape network (Canfield, 2003; Eaton et al., 2001; Gahtan et al., 2002; Weiss et al., 2006). Whether neural circuits subserving low and high threshold evasive behaviors are different or at least partially overlapping awaits investigation.

## Competing interests

We have no competing interests.

## Author contributions

VM and MBA contributed to conception of the study. SOC and VM participated in design of the experiments, carried out experiments and performed data analysis. NM performed experiments. SOC, MBA and VM contributed to drafting and revising the manuscript and all authors approved the final version of the manuscript.

## Funding

This work was supported by ANPCyT (PICT 20121578), Universidad de Buenos Aires (UBACyT 20020130300008BA), and CONICET (PIP2014 GI 11220130100729CO01) (VM).

## Acknowledgements

We thank Dr. Lidia Szczupak, Dr. Heike Neumeister and Dr. Thomas Preuss for discussion and for critically reading earlier versions of this manuscript. We also thank Ángel Vidal for invaluable technical assistance.

